# Dynamic Regulation of CD300b, and CD300f on Myeloid Cells in Oral Immunotherapy

**DOI:** 10.64898/2026.01.28.702205

**Authors:** N Epstein Rigbi, M Itan, S Avlas, Eytan Damari, L Nachshon, Y Koren, M Levy, M.R Goldberg, A Elizur, A Munitz

## Abstract

**Introduction:** Oral immunotherapy (OIT) induces desensitization in IgE-mediated food allergy, yet the role of myeloid cells in acquisition of tolerance is unclear. CD300 receptors regulate activation of myeloid cells, with CD300f acting as an inhibitory receptor and CD300b as an activating receptor. Their modulation during OIT may reflect effector cell reprogramming and serve as biomarkers of treatment response.

**Methods:** Thirty-five patients undergoing OIT were prospectively enrolled. Peripheral blood was collected at baseline and first 3 months of up-dosing; 19 patients completed sampling upon reaching maintenance. CD300b and CD300f expression in eosinophils, monocytes, and neutrophils was analyzed by flow cytometry. Allergen-specific IgE and IgG4 were measured by ImmunoCAP. Associations with clinical parameters were assessed using logistic regression. **Results**. Baseline CD300b was higher in patients with lower starting doses (*p*≤0.05). CD300f expression was lower in those with atopic dermatitis or multiple food allergies (*p*≤0.03). A significant downregulation of CD300b expression in the surface of eosinophils, monocytes, and neutrophils, was noted early in treatment (p≤0.05). Expression of CD300f was slightly but non-significantly increased. Longitudinally, the expression of CD300f increased on eosinophils, whereas CD300b expression decreased on the surface of monocytes and neutrophils. Specific IgE reduction correlated with downregulation of CD300b expression in monocytes *(R*=0.51, *p*=0.02) and higher CD300f expression in eosinophils *(R*=-0.45, *p*=0.05).

**Conclusions:** Downregulation of CD300b and upregulation of CD300f during OIT suggests myeloid cell reprogramming toward a less inflammatory phenotype. These dynamic changes in expression and their correlation with serologic markers of desensitization, suggest CD300b and CD300f as candidate biomarkers for understanding and monitoring OIT response.

**Key Message:**

1. OIT is associated with downregulation of the activating receptor CD300b and upregulation of the inhibitory receptor CD300f on myeloid cells.
2. Myeloid cell reprogramming accompanies OIT, extending immune tolerance beyond adaptive regulatory mechanisms.
3. CD300b and CD300f expression dynamics correlate with clinical dosing and serologic markers of OIT response.

## Introduction

Oral immunotherapy (OIT) has emerged as a promising approach for the treatment of food allergy(1,2). Recent studies have demonstrated that OIT can induce desensitization and, in some patients, sustained unresponsiveness(3). The underlying immunological mechanisms driving this process have been primarily attributed to the induction of regulatory pathways, including increased frequencies of regulatory T cells and the upregulation of immunoregulatory cytokines such as IL-10 and TGF-β(4). However, the involvement of innate immune effector cells from the myeloid lineage including eosinophils, neutrophils, and monocytes remains poorly understood(5). Whether these innate immune cells are passively modulated by the tolerogenic immune environment or actively contribute to the induction and maintenance of tolerance remains an open question. This distinction is critical given the increasing appreciation of effector cell plasticity and their potential regulatory functions in allergic inflammation(6). A better understanding of the phenotypic and functional alterations in these cells during OIT may provide novel insights into the mechanisms governing tolerance(7).

Members of the CD300 receptor family are expressed on multiple immune cell types, including eosinophils, neutrophils, monocytes, and mast cells(8). Among them, CD300a and CD300f are inhibitory receptors(9–11), whereas CD300b is an activating receptor(12,13). These receptors recognize lipid ligands such as phosphatidylserine and ceramide(14), and their expression has been associated with modulation of cell activation(15), cytokine release(16), and survival(17), particularly in the context of type 2 immune responses(18). In eosinophils, CD300f engagement attenuates degranulation and cytokine secretion(15,19), while in monocytes and neutrophils, CD300b has been linked to enhanced responses to microbial products(12,13,20).

Expression of CD300 molecules is dynamically regulated by environmental stimuli and inflammatory mediators(21). Type 2-associated cytokines, most notably IL-5, IL-33, IL-4, and IL-13 have been shown to modulate CD300 expression, particularly on eosinophils(8,22). IL-5 promotes eosinophil differentiation and survival and induces CD300f expression(23). Similarly, IL-33 can enhance CD300b and CD300f expression in myeloid cells, potentially modifying their activation threshold(23). Furthermore, CD300f is required for optimal activation of mast cells and eosinophils by IL-4^15^. These findings demonstrate that CD300 molecules are not only markers of cell activation states but also have functional roles in shaping effector responses during allergic inflammation.

Despite their emerging significance in allergy, the expression and functional relevance of CD300 receptors during immunotherapy and specifically OIT remain largely unknown(24). Herein, we examined the expression of CD300a, CD300b, and CD300f on peripheral blood eosinophils, neutrophils, and monocytes in patients undergoing OIT for food allergy. We aimed to determine whether OIT alters the expression of these receptors on effector cells and whether such changes may reflect active immune modulation associated with tolerance acquisition. This study provides insight into the potential role of effector cells and their inhibitory or activating receptors in OIT and may uncover biomarkers indicative of treatment response and immunological remodeling.

## Methods

### Study design and participants

Patients undergoing oral immunotherapy for milk, egg, peanut, sesame or tree nut allergy in the Shamir Medical Center, Israel, between March 2021 to July 2023 were recruited prospectively. All patients were recruited before starting the OIT program. The study was approved by the hospital ethical committee. All adult patients or care givers for pediatric patients signed informed consent.

### Oral Immunotherapy program

The OIT program is individualized, performed in an ambulatory care setting. Patients are eligible to participate over the age of 4 years. IgE-mediated food allergy was defined by evidence of IgE sensitization, either by skin prick test (SPT) or specific IgE, together with a positive oral food challenge (OFC), or a clinical history of an allergic reaction following accidental ingestion in the year prior to the start of OIT. Patients with a history of anaphylaxis and asthma were not excluded from treatment but asthma had to be well controlled throughout treatment. The program began with a 2-to-3-day dose-escalation phase during which the dose eliciting a reaction in each patient was established, and the starting dose (SD) determined. Dose escalation during the second and third rounds was limited to a maximum of 4-fold. Treatment subsequently consisted of alternate cycles of up-dosing and home-treatment phases until the full desensitization target dose (7200, 3000, 4000 and 6000 mg for milk, peanut, sesame and tree nut, and egg protein, respectively) was tolerated. Patients unable to reach this dose that were able to consume a minimal dose of at least 300 mg of the allergen protein, were defined as partially desensitized. Patients who stopped treatment all together were considered as treatment failure. A similar treatment protocol was used for patients of all ages.

### Blood sample collection and processing

Blood was drawn at three time points: during the first inductions days of OIT treatment representing baseline expression, during the first 3 months of up-dosing and upon finishing the up-dosing phase and reaching the maintenance dose. For 35 patients, blood was obtained at baseline and during the first phase of up-dosing. Only 19 of these patients completed the up-dosing phase within the time limits of the study and therefore blood was drawn for them in the 3 intended time points.

### Blood work analysis and flow cytometry

Identification of peripheral blood eosinophils, monocytes and neutrophils was performed using flow cytometry using the following antibodies: anti-CD16, anti-Siglec-8 (Biolegend, San Diego, CA), anti-CD14 and anti-CD45 (Biogems, Westlake Village, CA), anti-CD300a, anti-CD300b, anti-CD300e, anti-CD300f, and anti-CD300c (R&D Systems, Minneapolis, MN). Isotype controls including Rat IgG2A and Rat IgG2B were obtained from R&D Systems, Minneapolis, MN. Secondary antibodies including Donkey anti-Goat AlexaFlour 647, Goat anti-mouse AlexaFlour 488 and goat anti-rabbit AlexaFlour 647 were purchased from Jackson Immunoresearch, West Grove, PA). At least 10,000 events were acquired and CD300 receptor expression was analyzed using Kaluze software (Becman Coulter) as described^15^. Briefly, antibody staining was compared to isotype control and mean fluorescent intensity (MFI) fold increase (FI) was calculated.

### Analysis of IgE and IgG4

Analysis of allergen- specific IgE and IgG4 in the serum of patients were conducted on samples that were obtained from each patient at: baseline, early up-dosing phase and after the completion of up-dosing and reaching maintenance. For each patient, measurements of allergen specific IgE and allergen specific IgG4 for the patient’s specific allergen treated in OIT were measured using ImmunoCAP^TM^ (Thermo Fisher Scientific, Uppsala, Sweden).

### Statistical analysis

Statistical analysis aimed to compare CD300 receptor expression levels over time and to evaluate differences between patients who completed OIT up-dosing and maintenance versus those who did not finish up-dosing.

Normality of expression measures was assessed using the Shapiro-Wilk test. As distributions deviated from normality, continuous variables were summarized using median and interquartile range (IQR), and categorical variables were described as counts and percentages. Non-parametric tests were utilized for group comparisons and for longitudinal evaluation.

Differences between groups were evaluated using the Mann-Whitney U test for continuous variables and the Fisher’s exact test for categorical variables. Subgroup analyses, including comparisons of patients with atopic dermatitis and those with multiple food allergies, were also performed using the Mann-Whitney U test.

Longitudinal changes in receptor expression across time points (baseline, up-dosing, and maintenance) were assessed using the Friedman test for repeated measures. Violin plots were used to illustrate changes in receptor expression over time.

Associations of CD300 receptor expression with serological markers of immune modulation (allergen-specific IgE and IgG4) was analyzed using Spearman’s rank correlation coefficients (R), with results visualized using scatterplots and fitted trendlines.

The sample size was determined by the number of patients recruited during the study period and was not powered for formal hypothesis testing. Missing data were not imputed, and analyses were based on available data only.

A two-sided p-value <0.05 was considered statistically significant. All analyses were conducted using R version 4.3.2 (R Foundation for Statistical Computing, Vienna, Austria).

## Results

### Patient demographic and clinical characteristics

35 patients were recruited. All 35 completed blood work before the start of OIT and during the first 3 months of up-dosing. 19 patients completed the full up-dosing period and were able to reach full desensitization within the study period. Among the remaining 16 patients, 5 dropped out, one due to relapse of eosinophilic esophagitis and the rest due to emotional difficulties. A sub-analysis comparing clinical and demographic data between the patients who completed up-dosing (n=19) and those who did not (n=16) revealed a similar background (**Table S1**). The median age was 13 years (IQR 9, 19), 19 patients (54.3%) were female. Most patients had an atopic background, 18 patients (51.4%) had multiple food allergies (FA), 18 (51.4%) had asthma, 24 (68.6%) had house dust mite sensitivity and 17 (48.6%) had atopic dermatitis. Clinically, most patients had had a prior anaphylactic reaction (62.9%) with 17 (48.6%) requiring epinephrine injection. Four patients (11.4%) had undergone prior OIT for a different food allergen (**Table 1**).

**Table 1.** Patient demographic and clinical data.

| Category | Parameter |  | Patients<br>n=35 |
| --- | --- | --- | --- |
| Demographics | Age (years) |  | 13 (9,19) |
|  | Female gender |  | 19 (54.3%) |
| Atopic<br>background | Multiple food allergy |  | 18 (51.4%) |
|  | Asthma |  | 18 (51.4%) |
|  | HDM sensitivity |  | 24 (68.6%) |
|  | Atopic dermatitis |  | 17 (48.6%) |
| Clinical<br>Background | Prior anaphylaxis |  | 22 (62.9%) |
|  | Prior epinephrine injection |  | 17 (48.6%) |
|  | Prior OIT treatment (different allergen) |  | 4 (11.4%) |
| Oral<br>immunotherapy | Allergen treated | Milk | 9 (25.7%) |
|  |  | Peanut | 10 (28.6%) |
|  |  | Sesame | 8 (22.9%) |
|  |  | Walnut | 6 (17.1%) |
|  |  | Egg | 2 (5.7%) |
|  | Skin Prick Test (mm) |  | 7 (5,9) |
|  | Starting dose (mg protein) |  | 20 (5,40) |
|  | Interval 1<sup>st</sup> to 2<sup>nd</sup> sample (months) |  | 1 (1,3) |
|  | Duration of up-dosing (months), n=19 |  | 14 (11,16) |

Categorical variables presented as numbers and percentage, continuous variables presented as median and interquartile ranges.
HDM= house dust mite; OIT= oral immunotherapy

Patients underwent OIT for milk (25.7%), peanut (28.6%), sesame (22.9%), walnut (17.1%) and egg (5.7%). The median skin prick test was 7mm (IQR 5,9). The median starting dose was 20mg of protein (IQR 5,40) (**Table 1**).

### Early OIT Up-Dosing is Associated with Downregulation of CD300b and Mild CD300f Upregulation

To explore whether OIT modulates CD300 receptor expression, we analyzed the expression of CD300a, CD300b, CD300c, CD300e, and CD300f on eosinophils, monocytes, and neutrophils in 35 patients at baseline and after the early up-dosing phase. A sub-analysis comparing baseline expression levels in patients who eventually completed OIT (n=19) versus those still up-dosing or who dropped out (n=16) revealed no significant differences (**Table S2**), suggesting that baseline CD300 expression does not predict clinical trajectory. During early up-dosing, CD300b expression was significantly reduced across all three myeloid populations. Specifically, expression in eosinophils declined from a median fold increase in MFI of 1.94 to 1.6 (*p*=0.0437), on monocytes from 2.84 to 1.87 (*p*=0.0004), and on neutrophils from 2.5 to 1.69 (*p*=0.0085) (**Figure 1A-C**). This consistent downregulation may reflect early shifts in effector cell phenotype during immune modulation.

**Figure 1.**
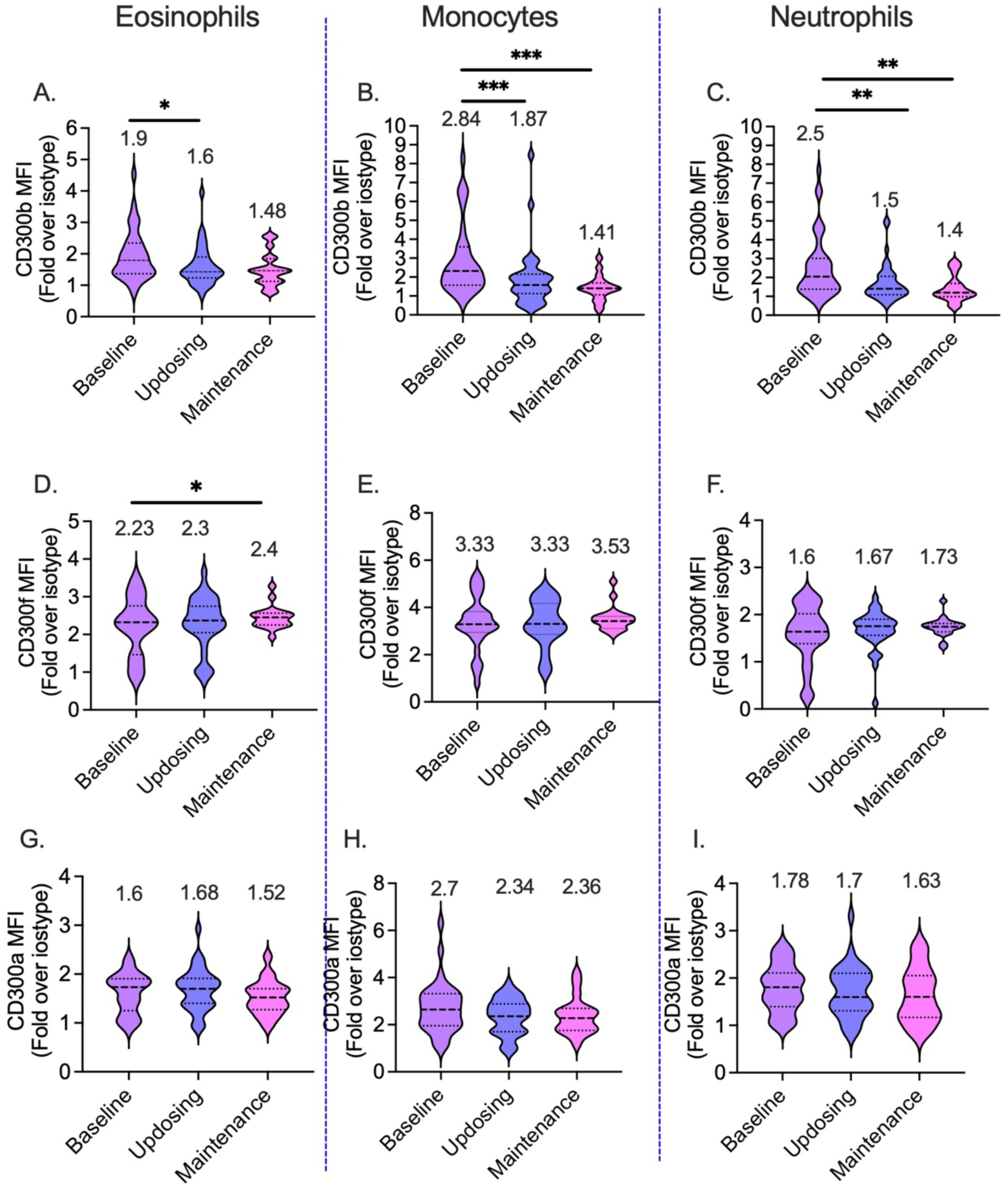
Early OIT up-dosing is associated with downregulation of CD300b on peripheral myeloid cells. Flow cytometric analysis of CD300 receptor expression in eosinophils (A, D, G), monocytes (B, E, H), and neutrophils (C, F, I) at baseline and during early up-dosing (n=35). CD300b expression was significantly reduced across all three cell populations (A-C), whereas CD300a and CD300f expression remained largely unchanged (D-I). Data is presented as fold increase (FI) in mean fluorescence intensity relative to isotype control. Paired comparisons were analyzed using the Wilcoxon signed-rank test; n=35, *-p<0.05, **-p<0.001, ***-p<0.0001.

While CD300a and CD300f were readily detectable on the surface of eosinophils, monocytes and neutrophils, their overall expression did not change (**Figure 1B-I**).

CD300c and CD300e were not detected on the surface of eosinophils, or neutrophils at either time point, with fluorescence intensities comparable to isotype controls (fold increase in MFI ∼1). Similarly, the expression of CD300c was undetected in monocytes. Nonetheless, CD300e was marginally expressed in monocytes under baseline conditions (fold increase in MFI 2.3) but its expression was not unchanged throughout the treatment regimen (Data not shown).

### CD300b and CD300f Expression at Baseline Reflects Starting Dose, Atopic Dermatitis, and multiple food allergy

Next, we examined whether baseline receptor expression is associated with any clinical parameters or treatment protocols. Baseline CD300b expression was significantly higher in patients who began OIT with a starting dose <20 mg protein (n=12) compared to those who started at ≥20 mg (n=7). For example, in eosinophils, the median fold increase (FI) of CD300b MFI was 1.85 vs. 1.37 (*p*=0.05); in monocytes the median FI of CD300b MFI was 2.46 vs. 1.86 (*p*=0.001); and in neutrophils the median FI of CD300b MFI was 2.1 vs. 1.63 (*p*=0.005) (**Figure 2A-C**). Notably, the expression of CD300b was decreased following OIT treatment only in the patient group whose starting dose was ≤20mg (**Figure 2A-C**).

**Figure 2.**
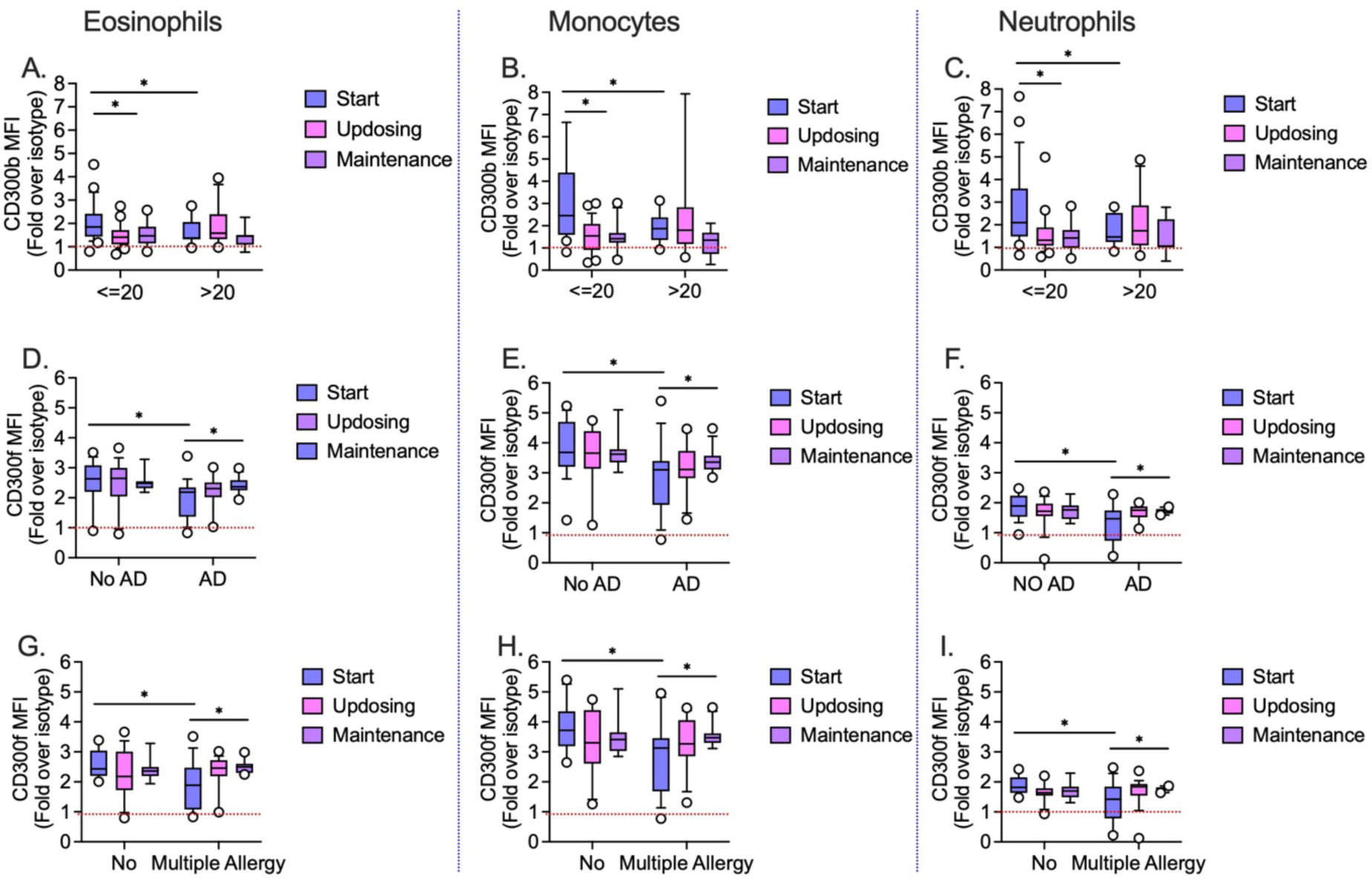
Baseline CD300b and CD300f expression is associated with OIT starting dose, atopic dermatitis, and multiple food allergy. Baseline CD300b expression in eosinophils, monocytes, and neutrophils stratified by starting dose (<20 mg vs ≥20 mg protein) (A-C). Baseline CD300f expression stratified by presence of atopic dermatitis (D-F) and multiple food allergy (G-I). Longitudinal changes demonstrate increased CD300f expression during OIT in patients with atopic dermatitis or multiple food allergies. Data is presented as median with interquartile range. Group comparisons were performed using the Mann-Whitney U test; n=35, *-p<0.05.

In contrast to CD300b, baseline expression of CD300f was lower on the surface of myeloid cells obtained patients with atopic dermatitis (n=11) and from patients with multiple food allergies (n=10) (**Figure 2D-I**). For instance, in eosinophils, the median FI of CD300f MFI was 2.18 in AD patients vs. 2.63 in non-AD patients (*p*=0.02); in monocytes the median FI of CD300f MFI was 3.1 in AD patients vs. 3.68 in non-AD patients (*p*=0.01); and in neutrophils the median FI of CD300f MFI was 1.46 in AD patients vs. 1.89 in non-AD patients(*p*=0.03) (Figure 2D-F). Similarly, patients with multiple food allergies (n=10) had lower baseline CD300f expression compared to those allergic to a single food (n=9, 1.88 vs. 2.43 in eosinophils (*p*=0.02); 3.12 vs. 3.71 in monocytes (*p*<0.01); and 1.42 vs. 1.81 in neutrophils (*p*<0.01) (**Figure 2G-I**). Importantly, assessment of CD300f expression during OIT treatment revealed that the expression of CD300f was increased during OIT treatment (from start to maintenance) in eosinophils, monocytes and neutrophils of patients with AD or multiple food allergies (**Figure 2D-I**).

Other clinical parameters including age, gender, skin prick test size, asthma status, prior OIT exposure, and baseline absolute eosinophil count (AEC) were not associated with differences in CD300 expression.

### Longitudinal Changes in CD300f and CD300b during OIT up-dosing

Nineteen patients completed longitudinal profiling of CD300 receptor expression across three time points (i.e., baseline, up dosing and maintenance). Among the receptors analyzed dynamic changes were observed in CD300b and CD300f in several cell types over the course of OIT up-dosing. Expression of CD300b in monocytes markedly declined over time, from a fold expression of 2.84 MFI at baseline to 1.41 MFI at the end of up-dosing (*p*<0.001). Similarly, the expression of CD300b in neutrophils, decreased from a baseline fold expression of 2.5 MFI to 1.4 MFI (*p*=0.006) (**Figure 1B-C**). Expression of CD300f in eosinophils increased significantly from baseline to the end of up-dosing (2.22 to 2.4, *p*=0.048), suggesting gradual upregulation during treatment (**Figure 1D**).

No additional longitudinal changes were detected in the expression of other CD300 family members on eosinophils, monocytes, or neutrophils during the same period.

### CD300 Expression Dynamics Correlate with Allergen-Specific IgE and IgG4 Responses During OIT

We further examined whether changes in CD300 receptor expression were associated with serological markers of immune modulation during OIT (n=19). Reduction in allergen-specific IgE between baseline and maintenance was associated with a larger decrease in CD300b expression in monocytes (*R*=0.51, *p*=0.022) (**Figure 3A**). A similar trend was observed in eosinophils as well(*R*=0.41, *p*=0.083) (b). No association was found between the decrease in CD300b expression and the increase in allergen-specific IgG4 levels over the course of treatment.

**Figure 3.**
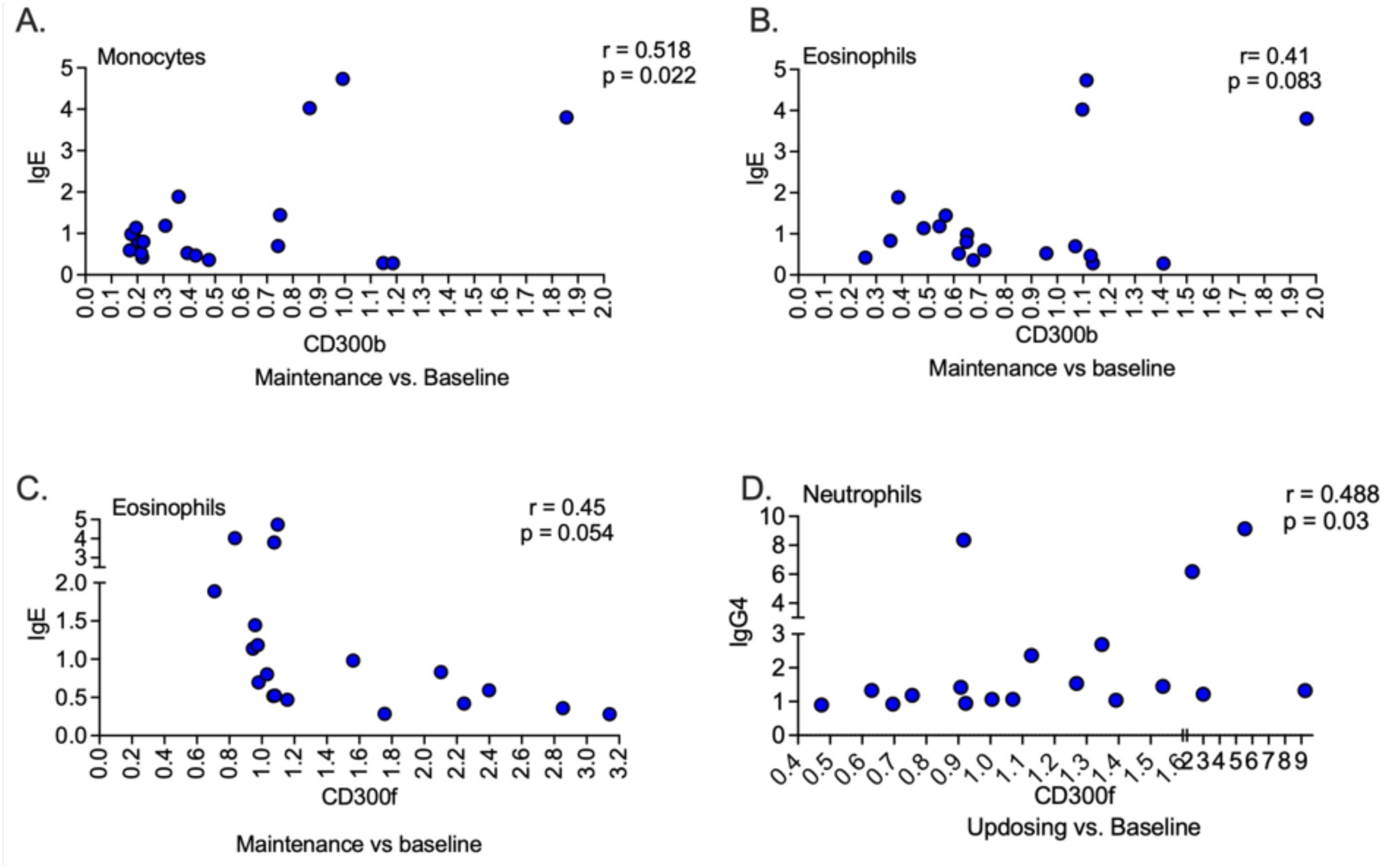
Changes in CD300 receptor expression correlate with allergen-specific IgE and IgG4 responses during OIT. Reduction in allergen-specific IgE from baseline to maintenance correlates with decreased CD300b expression on monocytes (A) and eosinophils (B). Higher CD300f expression on eosinophils correlates with greater IgE reduction (C), while increased CD300f expression on neutrophils correlates with increased allergen-specific IgG4 levels (D). Associations were assessed using Spearman’s rank correlation n=19.

Patients which exhibited higher expression of CD300f in their eosinophils throughout OIT were associated with a more pronounced decrease in allergen-specific IgE (*R*=-0.45, *p*=0.054) (**Figure 3C**). This association was not observed in monocytes or neutrophils (not shown). In contrast, neutrophils with elevated CD300f expression between baseline and early up-dosing showed a positive correlation with increased allergen-specific IgG4 (*R*=0.48, *p*=0.03) (**Figure 3D**). No additional associations were observed in other cell types or at different time points (not shown).

## Discussion

Oral immunotherapy (OIT) has been shown to induce clinical desensitization and, in a subset of patients, sustained immune tolerance in IgE-mediated food allergy(25). While adaptive immune mechanisms particularly the induction of regulatory T cells and immunoregulatory cytokines have been extensively studied(4,26), the contribution of innate immune effector cells to this process remains incompletely defined. In this study, we examined the dynamic regulation of CD300 family receptors on peripheral myeloid cells during OIT and demonstrate that treatment is associated with coordinated modulation of activating and inhibitory signaling pathways in eosinophils, monocytes, and neutrophils. These findings suggest that desensitization during OIT extends beyond adaptive regulation and involves reprogramming of myeloid innate effector cell phenotypes.

The most consistent and robust observation in our study was the progressive downregulation of the activating receptor CD300b, particularly on monocytes, observed both early during up-dosing and longitudinally throughout OIT. CD300b is an immunoreceptor known to amplify inflammatory responses through cooperation with pattern-recognition receptors such as TLR4 and through recognition of phosphatidylserine on apoptotic cells and stressed tissues(12,13,20,26). Its downregulation during OIT suggests attenuation of monocyte activation potential and may reflect a shift toward a more restrained innate immune state. This interpretation aligns with emerging models of immune tolerance in which suppression of innate effector activation accompanies adaptive immune regulation(27,28). Importantly, baseline CD300b expression was higher in patients requiring lower starting doses, and its subsequent decline correlated with reductions in allergen-specific IgE, supporting its relevance not only as a mechanistic marker but also as a potential indicator of treatment trajectory.

In parallel with CD300b downregulation, we observed a modest but significant increase in the expression of the inhibitory receptor CD300f, most prominently on eosinophils during longitudinal follow-up. CD300f is a well-characterized inhibitory receptor that regulates eosinophil survival, cytokine secretion, and responsiveness to type 2 cytokines(9,11,19,28). Engagement of CD300f has been shown to dampen eosinophil effector functions and to restrain allergic inflammation in both human and murine systems(15,19,23). In our cohort, higher CD300f expression on eosinophils was associated with greater reductions in allergen-specific IgE, suggesting that enhanced inhibitory signaling in eosinophils accompanies successful immune modulation during OIT. These findings are consistent with experimental models demonstrating that CD300f agonism can suppress food allergic responses and anaphylaxis(29), and further support a functional role for eosinophil inhibitory pathways in induction of tolerance.

Baseline CD300f expression was lower in patients with atopic dermatitis and in those with multiple food allergies. This is important since these two clinical phenotypes are associated with more severe or persistent allergic disease(15). Reduced expression of inhibitory receptors in these populations may reflect a heightened baseline activation state of myeloid cells, potentially contributing to disease chronicity. Notably, CD300f expression increased during OIT in these subgroups, suggesting that immunotherapy can partially reverse this inhibitory deficit. These findings support the concept that OIT not only suppresses allergen-specific adaptive responses but also recalibrates innate immune inhibitory pathways in patients with more complex atopic disease.

The reciprocal regulation which we observed (i.e., downregulation of the activating receptor CD300b alongside upregulation of the inhibitory receptor CD300f across distinct myeloid subsets) suggests a coordinated mechanism of immune restraint. Such bidirectional modulation may raise activation thresholds and limit effector responses during allergen exposure, thereby contributing to clinical desensitization(29). This pattern is consistent with prior work demonstrating cooperative and sometimes opposing functions among CD300 family members in regulating mast cell and eosinophil activation(11,15). Moreover, genetic studies in mice indicate that disruption of CD300 inhibitory pathways alters susceptibility to anaphylaxis(11), further underscoring the importance of balanced CD300 signaling in allergic responses(11).

Beyond mechanistic insight, our findings highlight CD300b and CD300f as potential peripheral biomarkers of OIT response. CD300b expression reflected initial dosing sensitivity and tracked with IgE reduction, while CD300f expression correlated with favorable serologic outcomes, including IgE decline and IgG4 induction. Given that these changes were detectable in peripheral blood, monitoring CD300 receptor expression may provide a practical and accessible window into immunologic remodeling during OIT and could aid in patient stratification or treatment monitoring.

Several limitations of this study should be acknowledged. First, longitudinal analyses were limited to patients who completed all sampling time points within the study period, resulting in a modest sample size. While the observed trends were consistent and biologically plausible, larger cohorts will be required to validate these findings. Moreover, we did not compare the results to a control group with food allergy that did not undergo OIT. Second, our analyses were confined to peripheral blood, which may not fully capture local immune dynamics at mucosal sites such as the gastrointestinal tract or skin, where allergen exposure and tolerance induction occur(30,31). Third, functional assays were not performed to directly assess how CD300 modulation alters myeloid cell behavior during OIT. Although correlations with serologic markers support a functional role, causality remains to be established. Finally, given the inclusion of multiple allergens and individualized treatment protocols, future studies should explore whether CD300 dynamics are allergen-specific or represent a shared pathway of immune tolerance.

## Conclusion

This study demonstrates that oral immunotherapy is associated with dynamic and coordinated modulation of CD300 receptor expression on peripheral myeloid cells. The observed attenuation of activating signals alongside enhancement of inhibitory pathways supports a model in which innate effector cell reprogramming accompanies and potentially reinforces adaptive immune tolerance during OIT. These findings expand the current understanding of OIT mechanisms and identify CD300b and CD300f as candidate biomarkers and regulatory nodes in allergen immunotherapy.

## Conflicts of interest

none declared

## Sources of funding

Ariel Munitz is supported by the US-Israel Bi-national Science Foundation (grant no. 2023029), by the Israel Science Foundation (grant no. 542/20), the Israel Cancer Research Fund, the Israel Cancer Association Avraham Rotstein Donation, the Cancer Biology Research Center (TAU), and the Azrieli Foundation Canada-Israel. Dr. Goldberg is funded by a Kamea grant from the Ministry of Health, Israel.

## Acknowledgments

We would like to thank the large staff of the Shamir Medical Center Allergy Clinic in their daily work and efforts: nurses, dieticians, respiratory technicians, laboratory workers and secretaries. We would also like to thank Jonas Lindholm and Lars Mattsson from Thermo Fisher, Sweden, for performing the antibody essays. Finally, we would also like to thank our patients for participating in this study.

## Notes

### Competing Interest Statement

The authors have declared no competing interest.

